# Intensity- and frequency-specific effects of transcranial alternating current stimulation are explained by network dynamics

**DOI:** 10.1101/2023.05.19.541493

**Authors:** Z. Zhao, S. Shirinpour, H. Tran, M. Wischnewski, A. Opitz

**Affiliations:** Department of Biomedical Engineering, University of Minnesota, Minneapolis, MN, USA

## Abstract

Transcranial alternating current stimulation (tACS) can be used to non-invasively entrain neural activity, and thereby cause changes in local neural oscillatory power. Despite an increased use in cognitive and clinical neuroscience, the fundamental mechanisms of tACS are still not fully understood. Here, we develop a computational neuronal network model of two-compartment pyramidal neurons and inhibitory interneurons which mimic the local cortical circuits. We model tACS with electric field strengths that are achievable in human applications. We then simulate intrinsic network activity and measure neural entrainment to investigate how tACS modulates ongoing endogenous oscillations. First, we show that intensity-specific effects of tACS are non-linear. At low intensities (<0.3 mV/mm), tACS desynchronizes neural firing relative to the endogenous oscillations. At higher intensities (>0.3 mV/mm), neurons are entrained to the exogenous electric field. We then further explore the stimulation parameter space and find that entrainment of ongoing cortical oscillations also depends on frequency by following an Arnold tongue. Moreover, neuronal networks can amplify the tACS induced entrainment via excitation-inhibition balance. Our model shows that pyramidal neurons are directly entrained by the exogenous electric field and drive the inhibitory neurons. Our findings can thus provide a mechanistic framework for understanding the intensity- and frequency-specific effects of oscillating electric fields on neuronal networks. This is crucial for rational parameters selection for tACS in cognitive studies and clinical applications.

## Introduction

Transcranial alternating current stimulation (tACS) is a non-invasive neuromodulation method that can directly interact with brain oscillations (1). Besides an increasing number of basic research studies, tACS has recently been explored as a clinical tool for the treatment of psychiatric and neurological disorders (1). With its various adjustable parameters, in particular stimulation intensity and frequency, tACS is well-suited for the development of personalized protocols. However, currently there is an incomplete mechanistic understanding of intensity- and frequency-specific effects of tACS at the neuronal and network level. Specifically, the effects of stimulation intensity and frequency on different neuron types and their interactions have yet to be explored.

The low-intensity stimulation from tACS induces sub-threshold membrane polarization to neurons, thus modulating the spike timings (1,2). There is a strong evidence that tACS modulates brain rhythms via entraining single neuron activity, as well as endogenous network oscillations, shown by *in vitro* (3–8) and *in vivo* rodent studies (9–11). Furthermore, studies in non-human primates have demonstrated phase-locking of ongoing neuronal spiking to the imposed waveform at relatively low intensities ≤0.3–0.4 mV/mm (12–14). Increasing intensity yields stronger entrainment with a larger number of neurons reaching the threshold for entrainment (12). However, for weak stimulation intensities, neurons may also be de-entrained to the natural endogenous oscillation (15). The entrainment effects are typically frequency-dependent and increase for higher stimulation amplitudes (11). In the study by Huang et al. (11), tACS was applied to the ferret brain using a frequency range between ± 4 Hz of the animal’s endogenous alpha frequency. They found that the range of frequencies at which entrainment occurred for neurons expands as the strength of the stimulus increases, also known as Arnold tongue (11).

In humans, neurophysiological evidence of tACS effects is more indirect. Several studies have shown that tACS delivered at the alpha and beta frequency range can increase motor evoked potential (MEP) amplitudes, thus indicating an increase in corticospinal excitability of the primary motor cortex (16–19). Effects of tACS on cortical excitability, as well as perception and cognition are often specific to a particular frequency range (e.g., theta, alpha, or beta) (20–23). Further, various studies have demonstrated frequency range-specific increases in EEG power (24,25). These observations are in line with the hypothesis that EEG frequency ranges are functionally distinct. Crucially, some human studies pointed toward frequency-specificity within a frequency range (26–28), although these findings are not demonstrated consistently (29).

Despite a wealth of existing literature on the effects of tACS across a wide range of stimulation parameters, theories that can integrate findings across studies are scarce. Computational models are crucial to develop a mechanistic understanding of experimental work. Additionally, modeling makes it possible to explore a wider range of stimulation parameters than is practical for experimental studies. Single-cell models demonstrated that pyramidal neurons are more sensitive to the exogenous electric field than other cell types due to their complex and elongated morphology (30,31). However, isolated single pyramidal neurons cannot account for all tACS effects. Neuronal network models with interconnected inhibitory and excitatory neurons show that the applied oscillating electric field might cause network resonance in neuronal populations. (5,32–35). Thus, stimulation frequencies that are closer to the endogenous frequency are more effective to entrain the network. Single-compartment (point) neurons are commonly used in such network models, however, they are not ideal for direct electric field coupling because of the lack of distance over which the change in extracellular potential occurs. The use of two-compartment models (36,37) allows for the direct capture of the effects of electric field. Because of the length of the dendrite, membranes can be de- and hyperpolarized at opposing ends of the neuron, accounting for differences in depolarization caused by external electric fields.

Here, we develop a unique approach using a neuronal network with two-compartment neurons to investigate the effect of tACS. We study the relationship between tACS intensities and neural entrainment and observed de-entrainment to the intrinsic oscillation at low stimulation intensities followed by re-entrainment to the external oscillation at a different phase for higher intensities. We further demonstrate that tACS entrains neuronal population dynamics via network resonance. Additionally, our modeling results suggest that network effects are an important factor in tACS-induced entrainment, which arise from the excitation-inhibition balance. Our network model provides a parsimonious account of known experimental results and provides important insights into the effects of intensity and frequency of tACS, which can inform future experimental studies.

## Results

We connected 800 two-compartment pyramidal neurons (PY) and 200 fast-spiking interneurons (IN) in a 3D volume (Fig.1 A, see Methods). To mimic the intrinsic network activity, we applied Poisson input to all neurons. We modeled tACS with the electric field vector aligned along the vertical axis of the network. We simulated network activity for 4 minutes: a 2-min baseline period (no tACS) and a 2-min tACS period. We measured the local field potential (LFP) from a virtua electrode inside the network (Fig. 1A). We first analyzed network activity at the baseline without external electric field. The LFP signal during the 2-min baseline had a strong spectral peak at ∼10 Hz due to the endogenous network activity (Fig. 1B). PY had a firing rate of 11.68 ± 2.70 spikes/sec, while IN had a firing rate of 56.99 ± 17.38 spikes/sec (mean ± standard deviation) (Supplementary Fig. 1).

**Figure 1.**
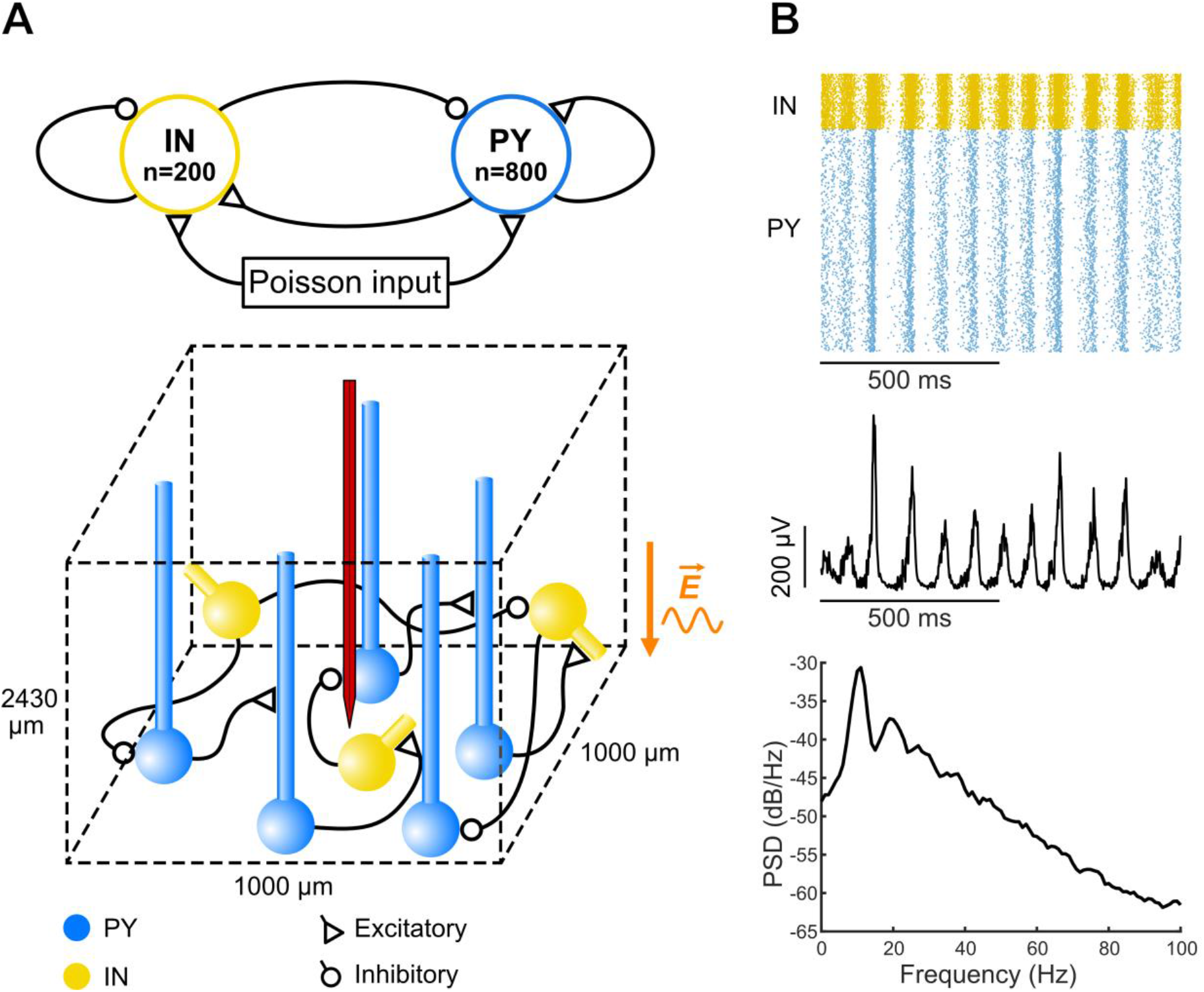
Network structure and endogenous alpha oscillation. **(A)** Top panel: a network of PY (n = 800) and IN (n = 200) population with excitatory and inhibitory synaptic connections within and between populations. Poisson input was applied to all neurons to generate ongoing network oscillation. Bottom panel: a schematic of the network in a 3D space. PY are orientated along the vertical axis while IN have random orientation to mimic the distribution of cortical neurons. LFP was measured in the extracellular space via a virtual recording electrode, shown in red. An alternating electric field (sinusoidal waveform) is applied to both PY and IN along the vertical axis. **(B)** Endogenous oscillation of the neuronal network. Top and middle panel: raster plot of neuron spiking activity and corresponding LFP signals for 1 simulated second. Bottom panel: power spectral density (PSD) of the LFP signals shows a peak at alpha range (10 Hz).

We then quantified the effect of 10Hz tACS on neural entrainment in the network. Trajectories on the polar plots in Fig. 2A show the effects of increasing electric field strength on spike timing of both PY and IN. Fig. 2B demonstrates a variability in PLV and phase preference of all neurons. In the baseline condition, all neurons were strongly entrained to the network’s 10 Hz LFP component. For an electric field intensity of 0.3 mV/mm, the neural entrainment decreased relative to baseline. However, at higher strengths, the electric field reinstated and increased entrainment at a different preferred phase. This indicates that weak external electric field reduces entrainment to the endogenous oscillation, but at higher intensities, tACS overcomes and increases entrainment to the tACS waveform. Interestingly, the phase preference of neurons shifts as the exogenous electric field interacts with the endogenous oscillation.

**Figure 2.**
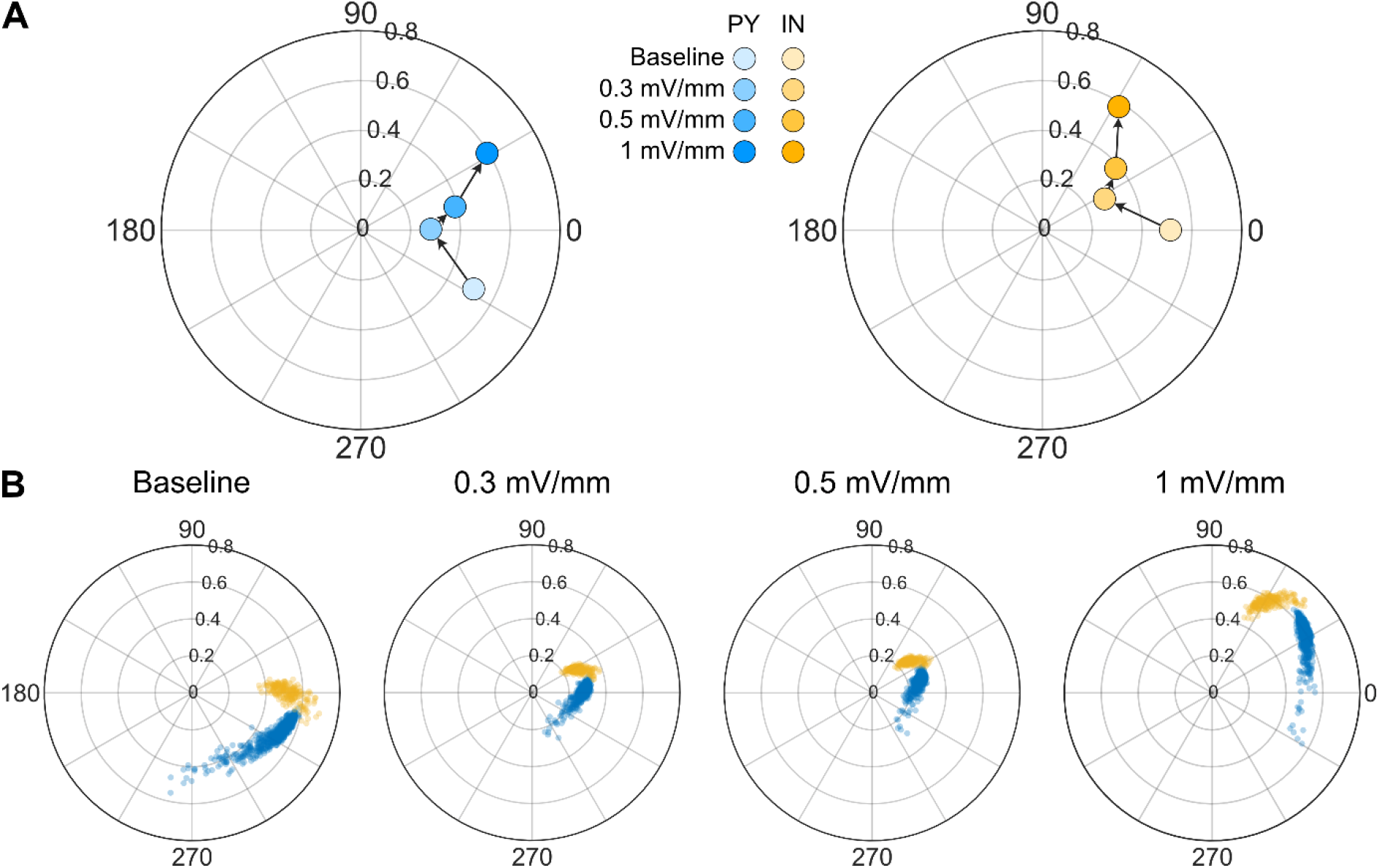
10 Hz tACS reduces entrainment and reinduces entrainment with phase shifts, as the stimulation intensity increases. Polar plots indicate the preferred phase (angle) and PLV (radial distance from the origin) of neurons in PY (blue) and IN (yellow). 0 degree corresponds to the peak of the oscillation and 90 degree corresponds to the falling edge of the oscillation. **(A)** Each dot shows the average across neurons in each population. Trajectories for Both PY and IN show reinstated entrainment **(B)** PLV and phase preferences of all neurons for four conditions in (A). Each dot represents an individual neuron.

Given that the network response was dependent on the stimulation intensities, we undertook a secondary analysis by stimulating the network model with a broad range of electric field intensities (between 0.1 and 1 mV/mm) and stimulation frequencies (between 1 and 30 Hz). For each simulation, we computed the firing rate and PLV of all neurons and created heat maps to visualize the evolution of the firing rate and PLV across simulations. Firing rates of both PY and IN neurons were not affected by weak electric fields (< 0.9 mV/mm) (Supplementary Fig 2). At higher electric field strength (> 0.9 mV/mm), the change in firing rate was less than 5%. For entrainment effects, we found high-entrainment regions centered on the endogenous alpha frequency (∼10 Hz) and first harmonic (∼20 Hz) for both PY and IN neurons (Fig. 3A). The range of frequencies at which entrainment occurred expands as the strength of the stimulus increases. Interestingly, we found that both neuron types show high baseline entrainment at around alpha and the first harmonic frequency. Starting from the baseline levels of entrainment, neurons were desynchronized relative to baseline as low intensity tACS was applied. At higher electric field strength, neurons were resynchronized to the external electric field, resulting in entrainment within the high-synchronization region. To illustrate this behavior, we discuss four specific cases of a PY’s entrainment within the stimulation parameter space (Fig. 3B). For 10 Hz tACS, as the electric field strength went up, the PLV first decreased from the baseline value and then increased. Polar histograms (Fig. 3B I-III) demonstrate a shift in the phases of the spike timings. At the same electric field strength (1 mV/mm), 10 Hz tACS entrained the neuron more effectively than 20 Hz tACS (Fig. 3B III and IV). To test if the synchronization regions also appear for different endogenous frequencies, we repeated the simulations for another endogenous cortical oscillation. We tuned our network model to exhibit an endogenous oscillation around the low beta band (14 Hz) (Supplementary Fig. 3). We observed similar high-synchronization regions on the PLV heat map (Supplementary Fig. 4). This demonstrates that the same qualitative behavior arises at different endogenous frequencies. These results suggest that cortical oscillations and tACS interact via phase locking at the single neuron and population levels.

**Figure 3.**
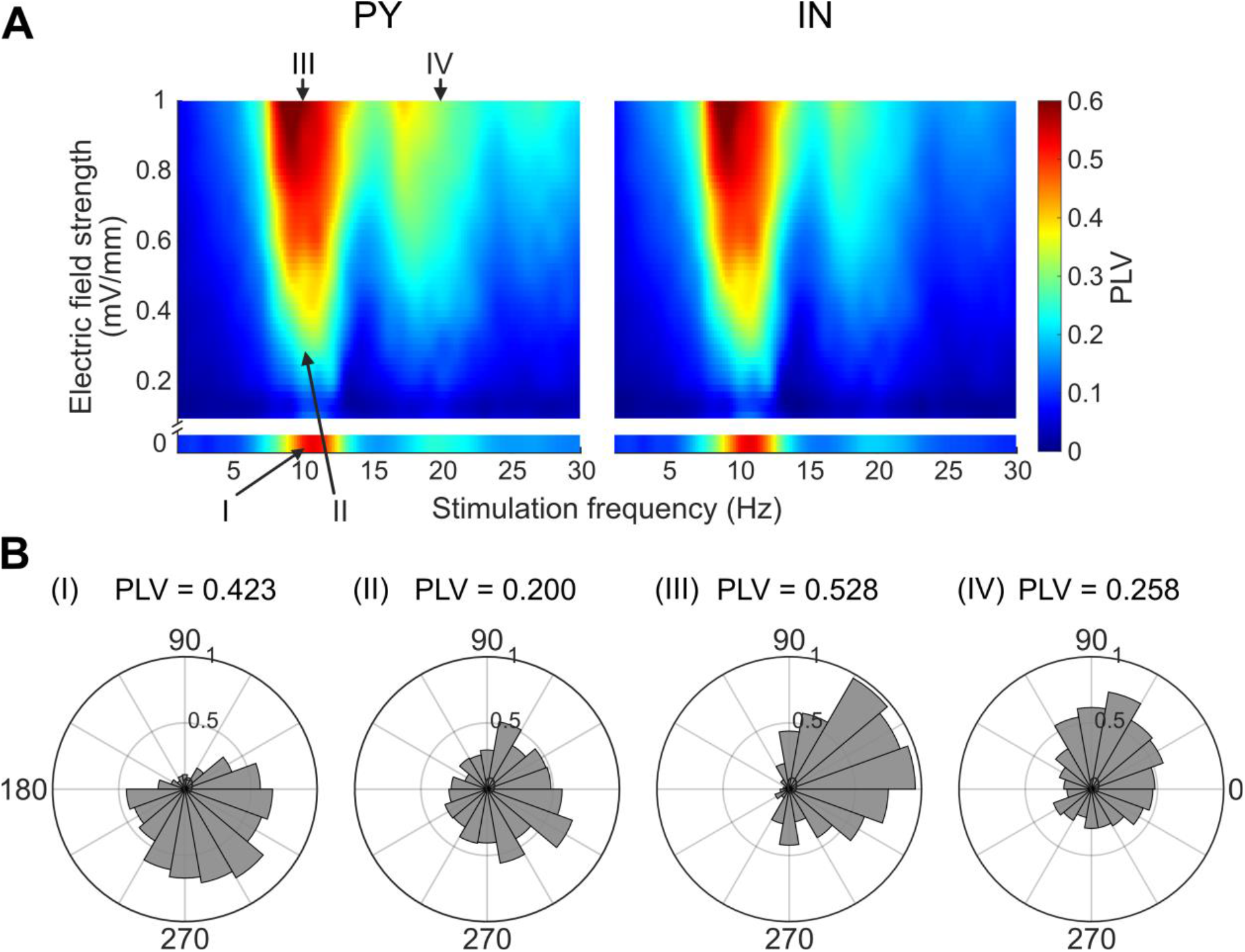
Network response to tACS with frequencies between 1 and 30 Hz. The lower section of each heat map represents PLV at baseline condition when no stimulation was applied. The upper section of each heat map shows intensities between 0.1 and 1 mV/mm. The color at each point shows the average across neurons. **(A)** Entrainment maps of neurons to tACS for each neuron type. Both PY and IN neurons show high PLV regions centered on the peak frequency (∼10Hz). **(B)** Four points on PLV map are selected to show different degrees of entrainment effects of a sample PY: (I) A point with high PLV under baseline condition. The sample PY is entrained to the endogenous 10 Hz LFP component. Rayleigh test. p-value = 1e-63. (II-IV) points with different stimulation parameters (II: 0.3 mV/mm and 10 Hz; III: 1 mV/mm and 10 Hz; IV: 1 mV/mm and 20 Hz. Rayleigh test. p-value = 1e-14, 1e-123, and 1e-26, respectively)

**Figure 4.**
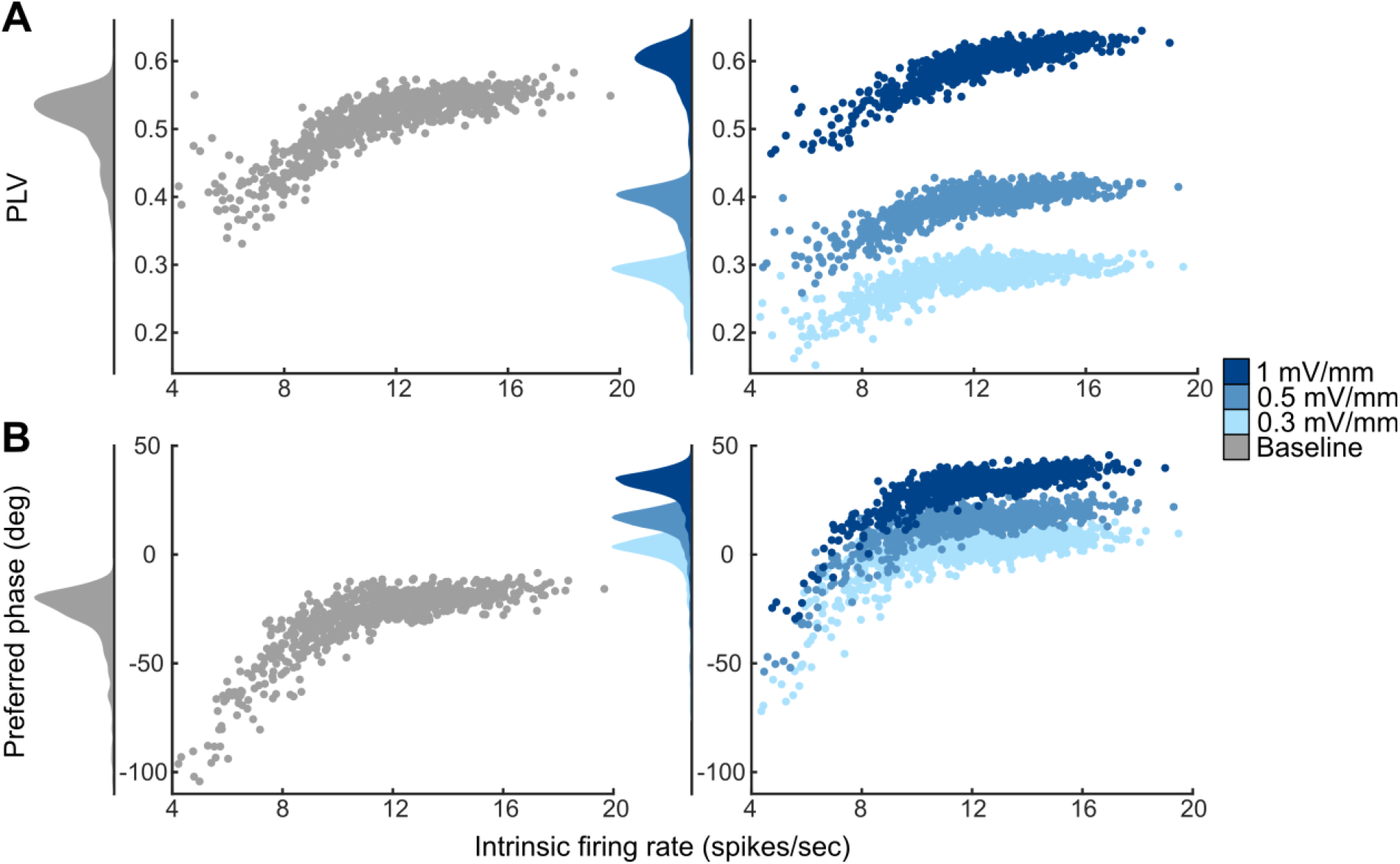
The effects of tACS differ between PY in the network. 10 Hz tACS was applied at three electric field strengths (0.3, 0.5, and 1 mV/mm). **(A)** Phase-locking values across all PY are shown for baseline and three electric field strengths. Individual dots indicate values for each PY in the network. PY that fire at higher intrinsic firing rates (*x* axis) show greater PLV. **(B)** Preferred phases across all PY. At baseline, PY show phase preferences before 0 degree (peak). PY with higher intrinsic firing rates tend to be entrained after the peak. Under tACS, the same trend remained. With increasing electric field strength, PY show larger backward phase shifts.

Since single neurons have a variable PLV and preferred phases under both baseline and tACS conditions across the population (Fig. 2B), we explored whether this spread of the PLV and phase preferences is due to the intrinsic firing rates of the neurons. We investigated the relationship between the neural entrainment of PY and their intrinsic firing rates under 10 Hz tACS, where we observed the largest tACS effects. Fig. 4 shows that most PY with higher intrinsic firing rates have higher PLV under both baseline and tACS conditions. Notably, this trend plateaus at higher firing rates (above 10 spikes/sec). Similarly, for phase preferences, PY with higher intrinsic firing rates are more likely to be entrained towards phases that are shifted backward (after peak) in time. Likewise, this trend plateaus after 10 spikes/sec. Under tACS, PY exhibit bigger phase shifts as electric field strength increases. Moreover, the logarithmic regression shows a high coefficient of correlation between PLV and firing rates (baseline condition: r = 0.859; tACS condition: 0.778, 0.822, and 0.892 for 0.3, 0.5, and 1 mV/mm respectively). In contrast, IN showed a different trend of PLV for varied intrinsic firing rates (Supplementary Fig. 5). IN with high firing rates were less entrained than IN with lower firing rates. Because our model neurons are interconnected within the network via excitatory and inhibitory synaptic connections, entrainment is affected not only by the difference between neuron firing rates and stimulation frequencies, but also by network effects arising from synaptic connections.

Finally, to study the mechanism of network effects on neural entrainment during tACS, we employed a simple network model with two neurons and one synaptic connection (Fig. 5). We investigated how the excitatory synapses (AMPA and NMDA) and inhibitory synapses (GABA_A_) affect the PLV of the postsynaptic neuron. For unbiased comparison, we included a control condition (no synapse) in which the postsynaptic neuron receives no synaptic connection from the presynaptic neuron while maintaining the same firing rates by adjusting the Poisson input. The excitatory or inhibitory synaptic weights were adjusted to keep the firing rates of PY and IN at 10 and 55 spikes/sec, respectively. Fig. 5 shows that the PLV of postsynaptic neuron was higher with an AMPA synapse than it was without synapse. Furthermore, the phase preference of the postsynaptic neuron was shifted further backward by NMDA synapses than by AMPA synapses (Supplementary Fig. 6). GABA_A_ synaptic connections, however, reduced the PLV of the postsynaptic neuron in comparison to the control condition (Fig. 5). Therefore, excitatory synapses enhanced the neural entrainment and caused phase shifts of the postsynaptic neuron. In contrast, inhibitory synapses reduced entrainment. Compared to the PLV of the isolated PY and IN, tACS effects on single neurons were amplified by network effects resulting from synchronized synaptic inputs. As a result, excitatory-inhibitory balance plays an important role in the network effects and the diversity of the neural entrainment in Fig. 4.

**Figure 5.**
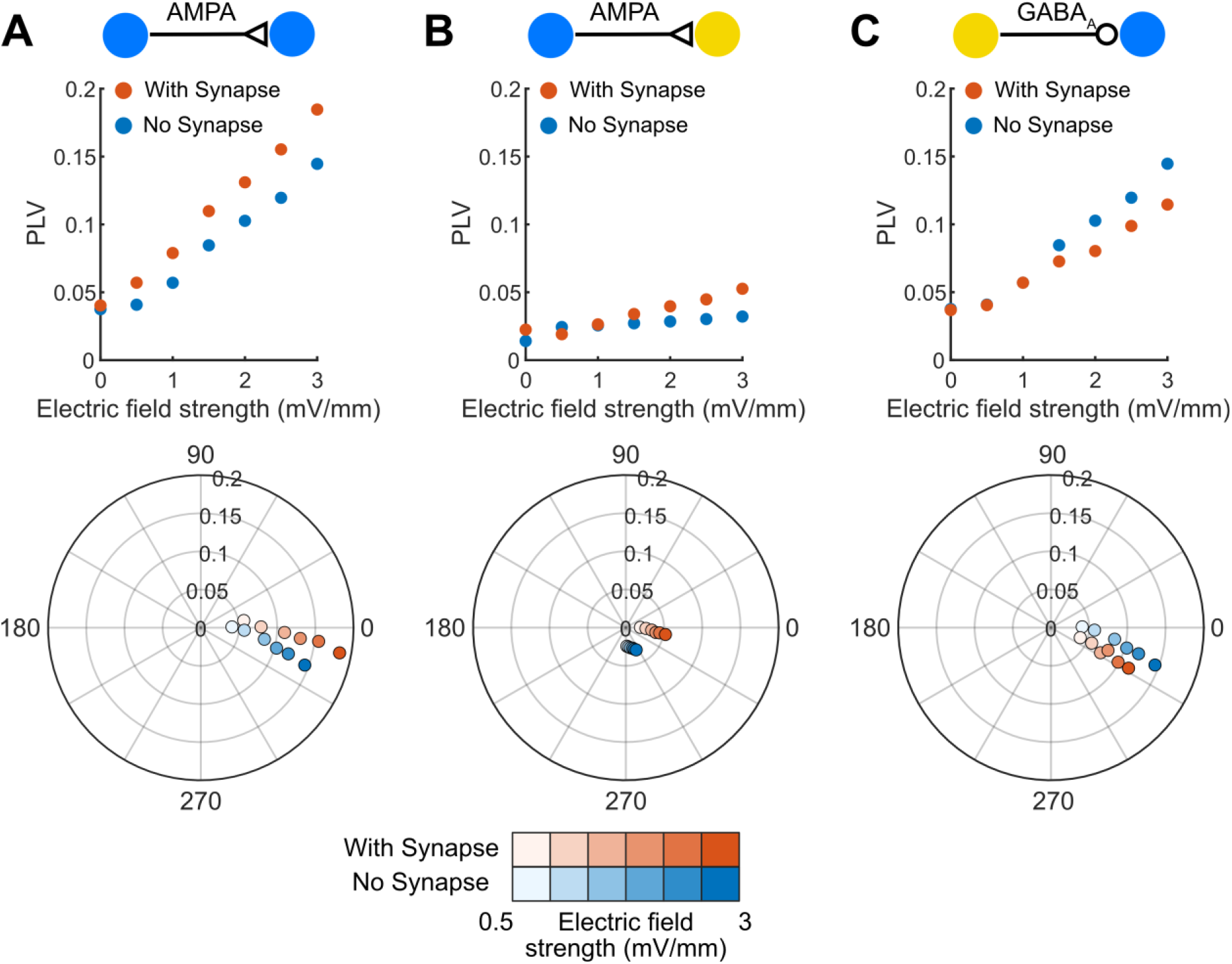
Synaptic connections affect entrainment of the postsynaptic neuron. Two neurons were connected through a monosynaptic connection. 10 Hz tACS was applied along the somato-dendritic axis of both PY and IN neurons. PLV plots and corresponding polar plots (the radial axis shows PLV) and show the effects of synapse on the entrainment of postsynaptic neuron (with synapse) relative to control (no synapse). **(A, B)** AMPA synaptic connection enhanced entrainment of postsynaptic neuron. **(C)** GABAA reduced entrainment of postsynaptic neuron.

Although both PY and IN were entrained by tACS at the frequency close to the endogenous frequency of the cortical oscillations (Fig. 3), we hypothesize that PY is the driving force of the network entrainment due to its higher sensitivity to the electric field. To determine whether the tACS-induced neural entrainment was driven by PY or IN, we performed a control analysis. Under 10 Hz tACS condition, we changed the electric field scaling factor (α) of each neuron type, which determines how neuron models respond to the external electric field. For each neuron type (PY and IN), we compared three conditions: 1) the neurons were not affected by electric field (α = 0), 2) the electric field coupling to neurons were doubled (α = 2), and 3) a default condition (α = 1). We computed PLV for all six conditions (Fig. 6). By doubling the α of PY, the entrainment of both PY and IN increased strongly (Fig. 6A). When PY were not affected by electric field (α = 0), the PLV of all neurons was lowered to less than 0.05, indicating that the network was not entrained by tACS. On the other hand, by changing the α of IN, the entrainment of both neuron types changed minimally, suggesting that IN neurons only marginally contribute to the direct entrainment of the network. This shows that PY were directly entrained by tACS, whereas IN indirectly contribute to the entrainment effects of exogenous electric field through synaptic inputs from PY.

**Figure 6.**
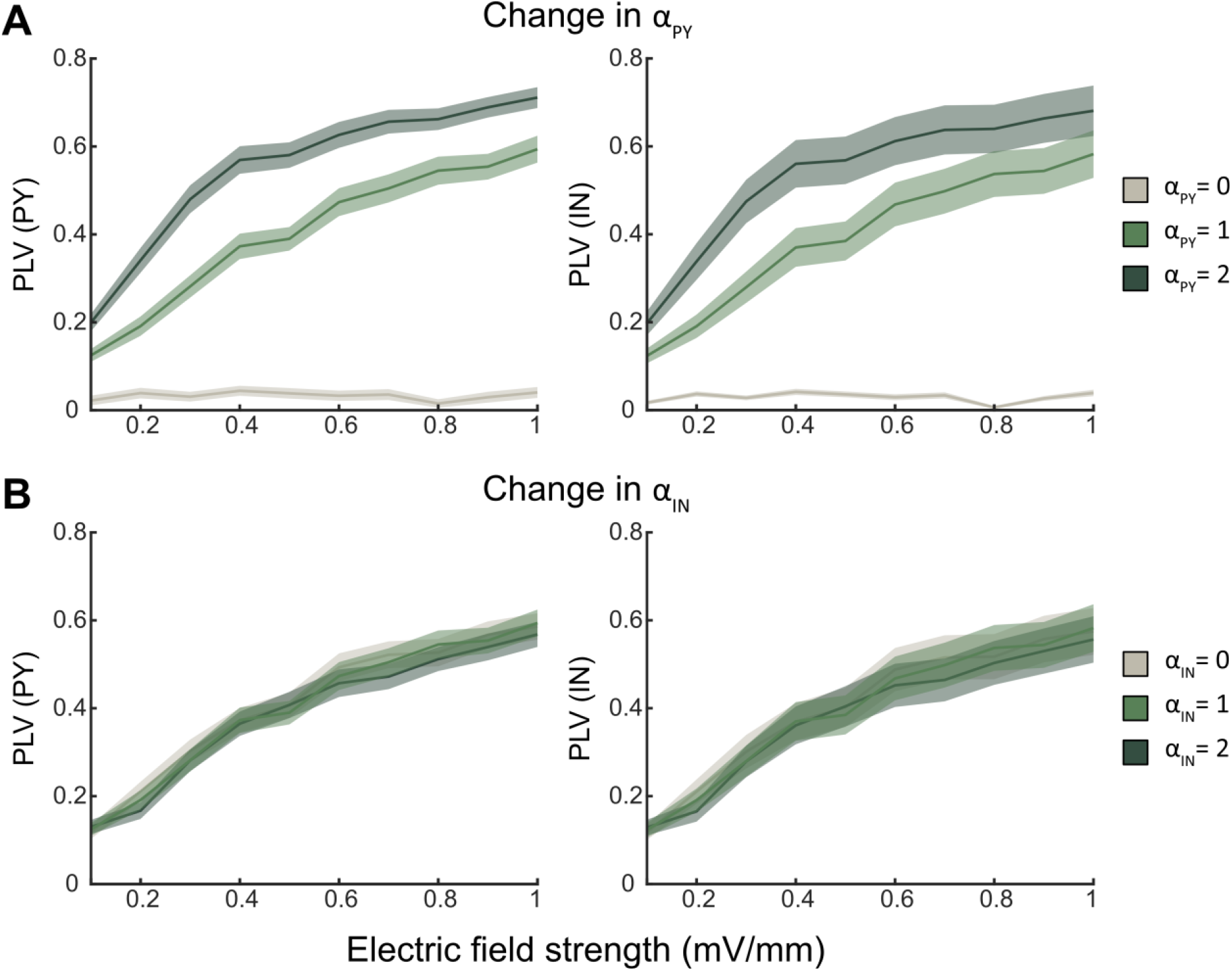
The entrainment of the network is affected by the sensitivity of neurons to external electric field. α value of 0 indicates that the model neurons are not affected by tACS; 1 indicates that the electric field sensitivity of neurons is the same as the original network; and 2 indicates that the sensitivity is doubled. **(A)** The sensitivity of all PY to the tACS (α_PY_) was altered in the network. The averaged PLV of all PY and all IN are shown (standard deviation is plotted as the shaded area). **(B)** The sensitivity of all IN to the tACS (α_IN_) was altered in the network. 10 Hz tACS was applied to the network in all conditions.

## Discussion

We have developed a computational neuronal network that consists of coupled two-compartment excitatory and inhibitory neuron models. Using this network model, we investigated how the intensity and frequency of tACS influence its interaction with ongoing cortical oscillations. We observed a non-linear relationship with first de-entrainment followed by re-entrainment at a different phase. The phase shift reflects the transition from endogenous entrainment to exogenous entrainment and was seen in both PY and IN. Besides intensity, we show that entrainment of ongoing cortical oscillations also depends on frequency. When the tACS frequency matches the endogenous frequency more closely, the phase-locking between the ongoing endogenous oscillations and the exogenous electric field increases. Additionally, network effects, resulting from excitation-inhibition balance, amplify the tACS-induced entrainment. We found that PY were entrained directly by the electric field and further drive the indirect entrainment of IN.

We observed that tACS entrains single neurons at intensities that are feasible for human studies, without causing considerable change in firing rates (Supplementary Fig 2). This in line with previous *in vivo* non-human primate studies (12,13). Moreover, the entrainment effects were non-linear. At intensities below 0.3 mV/mm, PLV decreased compared to baseline, suggesting desynchronization of neural firing in relation to endogenous oscillations. At intensities above 0.3 mV/mm, an increase in PLV was observed, which reflects synchronization of neural firing to the exogenous alternating current. In accordance, a recent meta-analysis suggested that AC stimulation can positively bias spike timing at intensities above 0.3-0.4 mV/mm (14). Further, our findings match a recent *in vivo* study in non-human primates where it was shown that transcranial alternating current stimulation (tACS) competes with the ongoing neural activity in controlling spike timing of single neurons (15). Depending on the stimulation intensity, this competition between tACS and ongoing neural activity determines the increase or decrease in the rhythmicity of neuron firing and shift of the preferred phase of spiking within an oscillatory cycle. Our results also demonstrated the same phenomenon. Entrainment to the ongoing oscillations was disturbed by exogenous electric field at low intensity. The network was then synchronized to the exogenous electric field at a different preferred phase at higher intensities (Fig. 2). This phase shift can be interpreted as the transition of phase-locking from the ongoing oscillation to the external electric field. Further, tACS induced phase shifts have recently been demonstrated in humans at a network level (38). Although these findings are preliminary, it shows that tACS-induced entrainment is non-linear and can involve changes in phase preference.

We explored the parameter space of tACS and found that it interacted with ongoing cortical oscillations through phase-locking mechanisms, as demonstrated by resonance zones in the stimulation parameter space centered around the network endogenous frequency (Fig. 3A and Supplementary Fig. 4B). This is referred to as an Arnold tongue, a synchronization region of coupled oscillators and an external force (39). This synchronization region has previously been demonstrated for a neural network model with applied external stimulation (5,11,32,34,35). It suggests that tACS can entrain network oscillations more effectively when stimulating with the endogenous frequency of the network oscillations (4,5,9). A recent study has shown that tACS entrains alpha oscillations by following the Arnold tongue in both ferret experiments and a computational model of the thalamo-cortical system (11). Currently no direct evidence, such as single unit recordings, of the Arnold tongue phenomenon in humans has been reported. However, some indirect evidence suggests that effects of tACS are more pronounced when using individual peak frequencies, compared to standard values (26) or control frequencies slightly above or below the peak frequency (27,28). It should, however, be noted that such strong frequency dependency is not universally observed (29).

We observed variability in PLV and phase preferences of neurons while the whole network was entrained (Fig. 2). This was further investigated based on the difference in intrinsic firing rates between neurons (Fig. 4). A computational modeling study (30) suggested that at the same stimulation frequency, an isolated single neuron that fires at higher rates relative to the stimulation frequency has lower PLV than the same neuron that fires at lower rates. This might explain why the PY trend plateaus after 10 spikes/sec during 10 Hz tACS and the PLV of IN shows an overall downward trend. Since our model neurons are coupled in the network via excitatory and inhibitory synaptic connections, the variability in PLV cannot be simply explained by the intrinsic firing rates of single neurons. Therefore, neural entrainment is affected by network dynamics.

Neuronal networks can amplify the tACS induced entrainment. Studies using *in vitro* and *in vivo* models provided evidence for the idea that active networks may be more sensitive to electric fields than isolated single cells (5,9,11). Our model also demonstrates that neuronal networks can amplify the tACS induced entrainment via excitation-inhibition balance. Excitation enhanced the neural entrainment via AMPA and NMDA synaptic connections to the postsynaptic neurons (Fig. 5). Recurrent feedback between excitatory and inhibitory neurons causes an increase in inhibition after excitation is increased. Such balance allows for generating stable periods of activity and modulating network oscillations (40,41). Previous research has suggested that tACS may target specific types of neurons: pyramidal neurons have been shown to be more susceptible to electric field while small interneurons are less susceptible (6,30,42). Other studies suggest that network interactions make tACS have especially strong effects on interneurons, which drives the remaining neurons (11). However, our modeling results show that, despite being highly entrained, interneurons were primarily entrained indirectly by the excitatory neurons (Fig. 6). This means that without connections from PY, IN do not highly synchronize to the exogenous current.

Our model, while capable of reproducing existing experimental findings and highlighting crucial tACS mechanisms, has some limitations. First, we did not include varied orientations and locations of pyramidal neurons. Although we only study tACS effects on a local network level, it can affect broad area in the cortex (38). Second, we did not consider various lengths of two-compartment PYs and INs. To generate more accurate estimates of the tACS effects while remaining computationally efficient, we tuned our cortical neuron models to match the PLV of those with realistic morphologies. However, different morphologies of neurons can potentially lead to a larger variability in their response to tACS (30,31,37). Third, we did not investigate tACS-induced modulation of neural plasticity mechanisms that could explain stimulation-related aftereffects (24,43,44). Recent study has shown that tACS-induced phase shifts are related to plasticity (38). Future work could incorporate spike-timing-dependent plasticity (STDP) into the network and investigate the NMDA receptor-mediated plasticity during tACS.

In conclusion, this study used computational modeling to provide insight into the mechanisms of tACS effects on neuron populations. The findings highlight the importance of tailoring stimulation parameters to specific intensities and frequencies based on the endogenous neural oscillation. Different stimulation intensities would result in diverse neural entrainment effects. On the other hand, using a stimulation frequency that matches the intrinsic neural oscillation, phase-locking can occur at lower electric field strengths. Our model allows the exploration of tACS parameters and could pave the way for more effective use of tACS in clinical settings.

## Materials and methods

### Neuron models

We implemented conductance-based two-compartment neuron models consisting of soma and a dendrite (Fig. 1A). Single-compartment neurons are commonly used in network modeling, however, two-compartment models can capture the effects of electric fields (36) while still being computationally efficient. We used NetPyNE (45), a Python package to simulate and model biological neuronal networks based on the NEURON simulation environment (46). The membrane potential of the model neuron can be calculated by the following equation based on the cable theory (47):

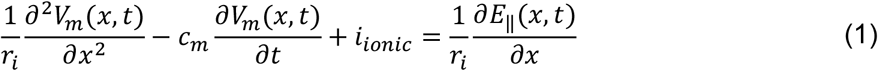

where *E*_||_*(x,t)* is the parallel component of the electric field along the neuron compartment, *V*_*m*_ is the membrane potential, *r*_*i*_ is the cytoplasmic resistance, *c*_*m*_is the cell membrane capacitance, *i*_*ionic*_ are the currents passing through membrane channels, *x* is the location of neuron compartment, and *t* is the time.

The morphologies of each type of neuron are shown in Fig. 1A. Large pyramidal neurons (PY) are often considered to be more susceptible to electric fields than small interneurons (IN) (30,31,48). Active currents in PY include a fast sodium current (I_Na_), fast potassium current (I_Kv_), slow non-inactivating potassium current (I_km_), leak current (I_L_), calcium current (I_Ca_), and Calcium-dependent potassium current (I_KCa_). The Hodgkin-Huxley-style kinetic equations of these currents are adapted from the model (49). Membrane electrical properties, such as membrane capacitance and ion channel conductance, were modified to generate a regular-spiking firing pattern. The IN model was adapted from the single-compartment model (50) and a dendritic compartment is added for electric field coupling. IN contain I_Na_ and I_Kv_ currents to create fast-spiking activity. The parameters used for the morphologies of the neurons are shown in Supplementary Table 1.

### Single neuron model tuning

The reduced neuron model with simple morphologies helps speed up the simulation of the large network model. Neuron models with realistic morphologies can produce accurate predictions of the electrophysiological effect. To address this, we tuned the two-compartment models so that their responses to the electric fields are similar to the realistic neurons (30). To this end, the electric field coupling to the model neuron was scaled to match the realistic neurons. We matched the entrainment of the reduced neuron model to the morphologically realistic neuron models by adjusting the effective length and neuron membrane’s electrical properties, such as the membrane capacitance and resistance.

### Network architecture

The network consists of 1000 neurons with 800 PY and 200 IN (51). PY in the network were aligned to the vertical axis, and IN were distributed in 3D space (1000 μm × 1000 μm × 2430 μm) with random orientations (Fig. 1A) to reflect the typical cytostructure of cortical columns. Neurons are connected to other neurons in- or outside their population. Excitatory and inhibitory synaptic connections within the network were constructed as shown in Fig. 1A. Excitatory synapses connect to the dendrites, and inhibitory synaptic connections on PY and IN are located on the soma (52). All neurons in the network received background Poisson input to allow for spontaneous firing to mimic the ongoing *in vivo* activity of neurons.

### Synaptic connections

The electric current I_syn_ that results from a presynaptic spike was modeled as the follows (53,54):

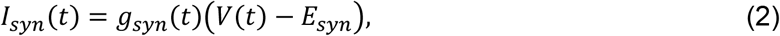

where the effect of transmitters binding to and opening of postsynaptic receptors is a conductance change, *gsyn*(*t*), in the postsynaptic membrane. *V*(*t*) denotes the transmembrane potential of the postsynaptic neuron and E_*syn*_ is the reversal potential of the ion channels that mediates the synaptic current. If the neuronal transmembrane potential is below the reversal potential, presynaptic spike arrival leads to a depolarization of the neuron. The equation is used to describe the time course of the synaptic conductance (53,54):

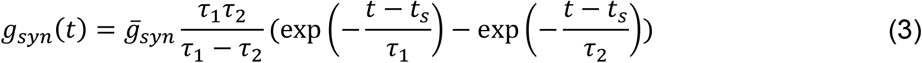

where *t*_*s*_ is the time of a presynaptic spike, *τ*_1_ and *τ*_2_ characterize the rise and decay time of the synaptic conductance in the dual exponential function. The conductance change resulting from α-amino-3hydroxy-5-isoxazolepropionic acid (AMPA), N-methylD-aspartate (NMDA), and γ-aminobutyric acid type A (GABA_A_) mediated synaptic current are simulated using a dual exponential function (Eq. 3). Parameters used for excitatory and inhibitory synaptic conductance are adapted from (55) and are shown in Supplementary Table 2. The synaptic conductance resulting from the background Poisson input is also modeled as a dual exponential decay with Esyn = 0 mV, τ_1_ = 2 ms and τ_2_ = 10 ms (30).

### *Modeling cortical* oscillations

To generate an ongoing, endogenous oscillation in the network, we adjusted the Poisson input to both the PY and IN, as well as the synaptic weights within and between the population. We sought to model a cortical oscillation, such as the alpha-band oscillations (8-13 Hz), which are dominant oscillations in the human brain (56), and have previously been effectively targeted in a number of tACS human studies (43,57,58). Additionally, to determine whether the same entrainment effects persist for various network dynamics, we modeled an endogenous oscillation at low beta frequency (14 Hz). We implemented a reciprocal local connectivity scheme (59). The parameters used for the network connectivity are shown in Supplementary Table 3. To add the heterogeneity to the network, we used a distinct seed of the Poisson input for each single neuron.

### Local field potential (LFP)

The LFP signal was obtained by summing the extracellular potential induced by each segment of each neuron at the electrode location. Extracellular potentials were calculated using the Line Source Approximation (60,61) and assuming an Ohmic medium with conductivity sigma = 0.3 mS/mm (62). The LFP of the network was computed and filtered using a 2^nd^ order Butterworth bandpass filter (cutoff frequency = 0.1 Hz and 300 Hz).

### Modeling tACS

To couple the tACS electric fields to the neuron model, quasipotential (*ψ*), was calculated for each neuron compartment based on the external electric field induced by tACS (63). When the electric field is assumed to be uniformly distributed, the quasipotential equation can be simplified as follows (64):

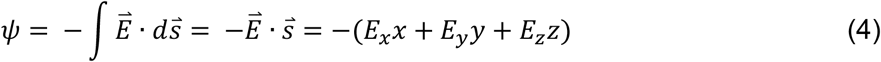

Where 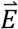 is the electric field vector, 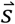 is the displacement. E_*x*_, E_*y*_ and E_*z*_ indicate the Cartesian components of the electric field in a three-dimensional space, and x, y, and z denote the Cartesian coordinates of each neuron compartment. The quasipotentials were calculated at each neuron segment and were set as the extracellular potentials in the “extracellular mechanism” in NEURON environment (42).

We used a sinusoidal tACS waveform and simulated a range of electric field strengths (0 to 1 mV/mm in increments of 0.1 mV/mm) and frequencies (1 to 30 Hz in increments of 1 Hz). Using this range, we cover typical intensities and frequencies applied in humans (14) as well as explore the effect of tACS at a higher dosing range.

### Phase-locking analysis

We quantified neural entrainment by calculating the phase-locking value (PLV), which measures spike timing synchronization with respect to an ongoing signal. The PLV is calculated as follows (65):

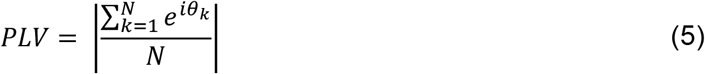

Where N is the number of spikes and θ_*k*_ denotes the phase of the oscillation at the time where the k^th^ spike occurs. A uniform distribution of spike timings over all phases (0 to 2π) results in a PLV of 0, while a value of 1 means perfect synchrony to a specific phase of tACS. Under baseline condition, we measured PLV with respect to the LFP signal at a frequency of interest (filtered into a ±1 Hz range). While for tACS condition, PLV was calculated with respect to the tACS waveform(15). Note that for generating heat maps, cubic interpolation was used to smooth over data points.

### Model simulation and Data analysis

The simulations were done in NEURON version 7.6 using a fixed-time-step cnexp method (modified Crank-Nicolson method) with a time increment dt = 0.05 ms. The total duration of an individual simulation was 4 minutes – a 2-min baseline period (no tACS) and a 2-min tACS period. All simulations were done using the Minnesota Supercomputing Institute (MSI). All post-simulation analyses were done in MATLAB 2021b.

## Acknowledgements

Research presented here was supported by the National Institute of Health (NIH grant: RF1MH124909). The authors acknowledge the Minnesota Supercomputing Institute (MSI) at the University of Minnesota for providing resources that contributed to the research results reported within this paper. URL: http://www.msi.umn.edu

