## Supplementary for "Intensity- and frequency-specific effects of transcranial alternating current stimulation are explained by network dynamics"

### Supplementary materials

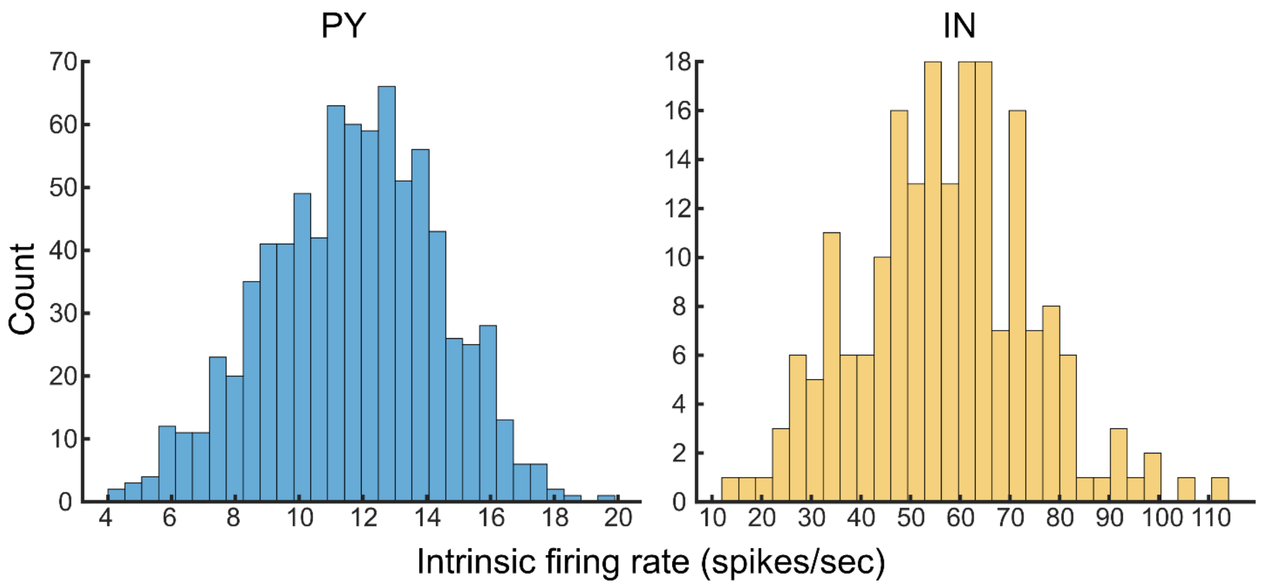


**Supplementary Figure 1**. **Firing rate distributions for PY and IN during alpha oscillation.** The firing rates are 11.68 ± 2.70 spikes/sec for PY and 56.99 ± 17.38 spikes/sec for IN (mean ± standard deviation).

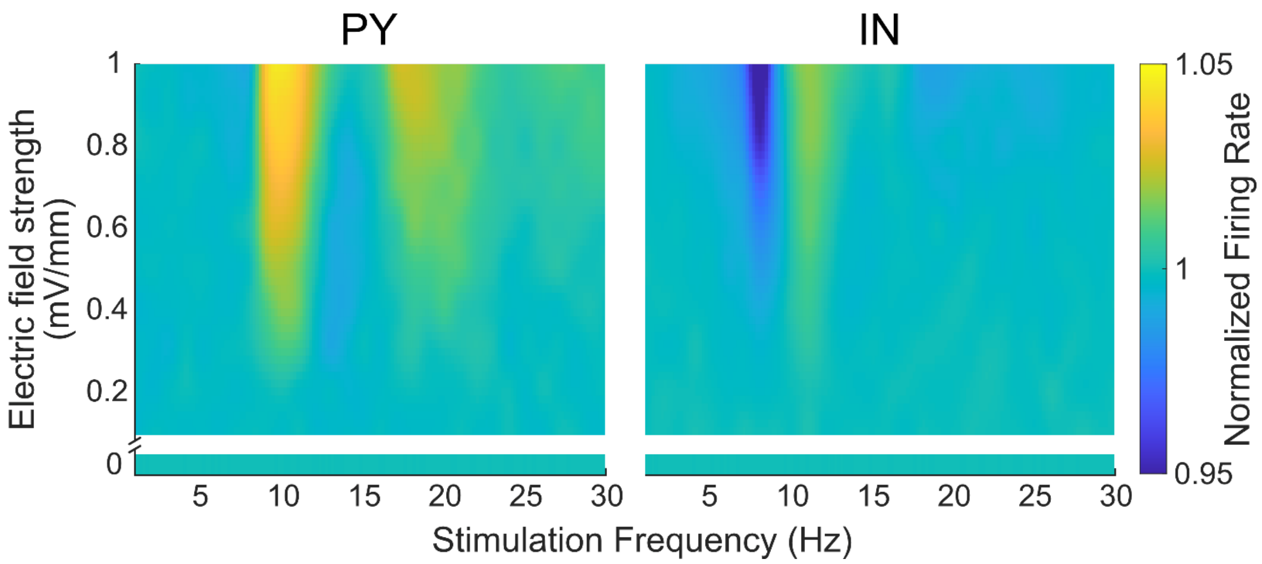


**Supplementary Figure 2. Normalized firing rate maps for each neuron type.** The network has an endogenous oscillation around 10 Hz. Firing rates of both PY and IN neurons were not affected by weak electric fields (< 0.9 mV/mm). At higher electric field strength (> 0.9 mV/mm), a change in firing rate was less than 5%.

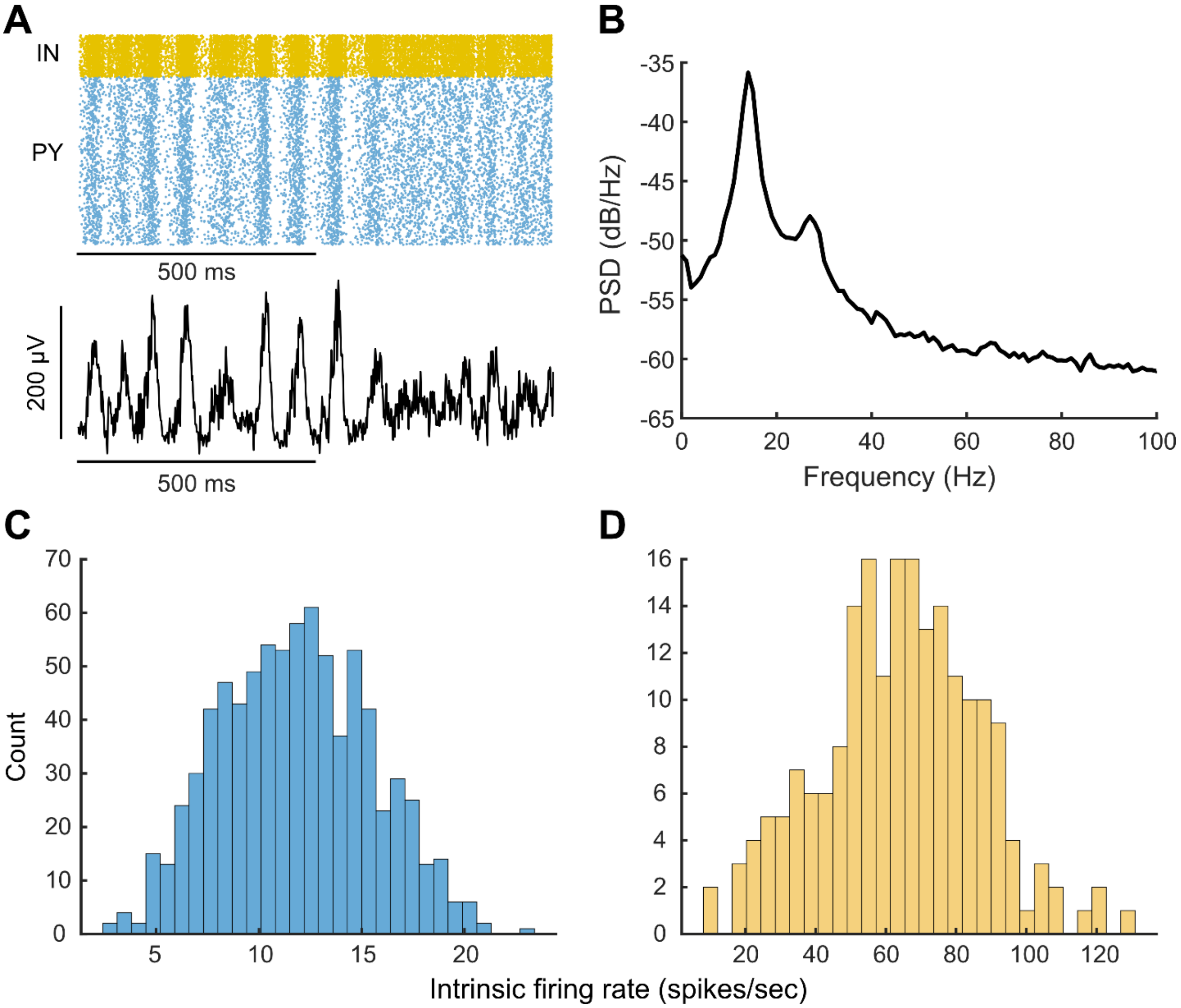


**Supplementary Figure 3. Endogenous low beta oscillation. (A)** Raster plot of neuron spiking activity and corresponding LFP signals for 1 simulated second. **(B)** The network oscillation shows a power peak at low beta range (14 Hz). **(C, D)** Firing rates of PY and IN were distributed in a wide range (PY: 11.80 ± 3.67 spikes/sec, IN: 64.06 ± 22.40 spikes/sec, mean ± standard deviation).

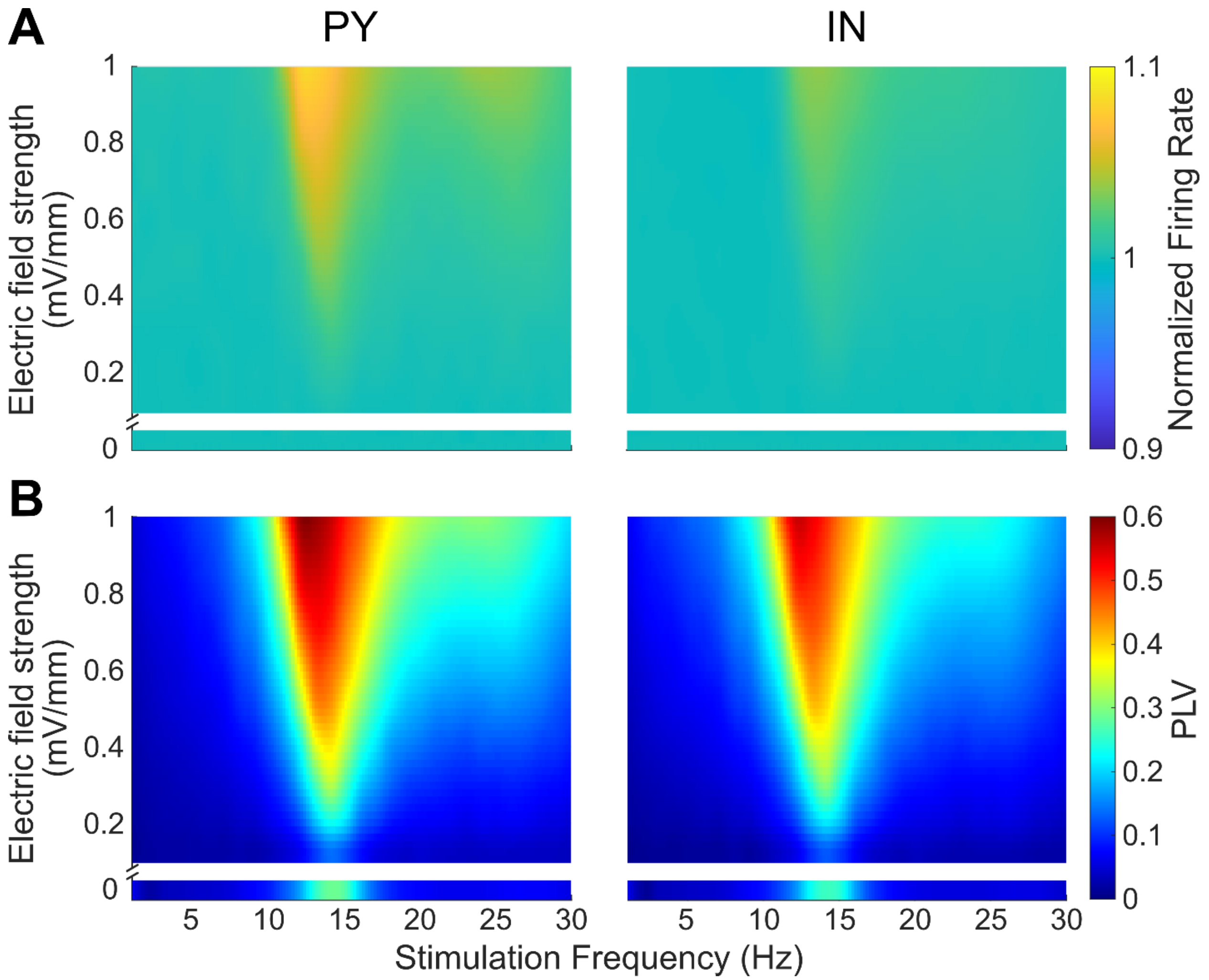


**Supplementary Figure 4. Network response to tACS with different frequencies and intensities.** The lower section of each heat map represents normalized firing rate and PLV at baseline condition when no stimulation was applied. The upper section of each heat map shows intensities between 0.1 and 1 mV/mm. The color at each point shows the average across neurons. **(A)** Normalized firing rate (change compared to baseline) maps for each neuron type. With 14 Hz tACS, firing rates of both PY and IN neurons were not affected by weak electric fields (< 0.6 mV/mm). At higher electric field strength (> 0.6 mV/mm), change in firing rate was less than 10%. **(B)** Entrainment maps of neurons to tACS for each neuron type. Both PY and IN neurons show high synchronization regions centered on the peak frequency (~14Hz).

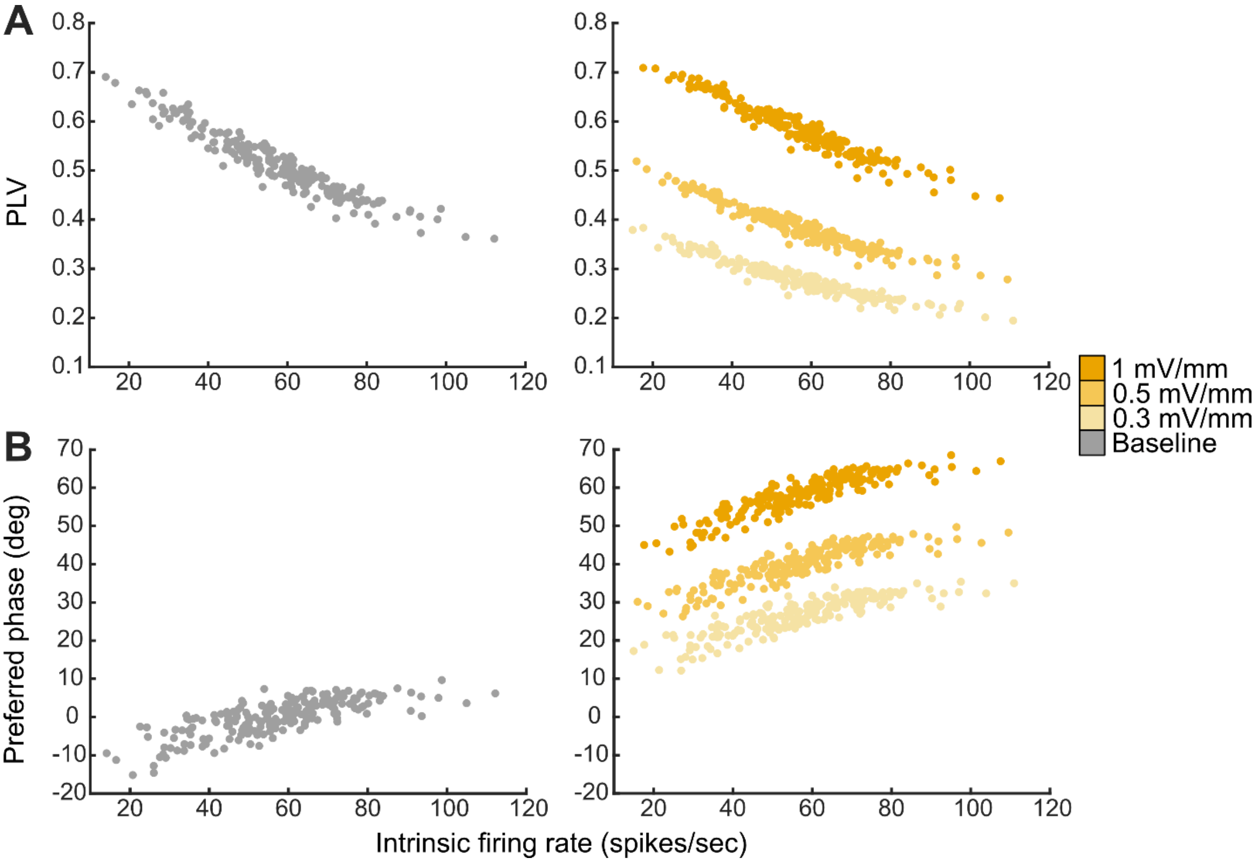


**Supplementary Figure 5. The effects of tACS differ between IN in the network.** 10 Hz tACS was applied at three electric field strengths (0.3, 0.5, and 1 mV/mm). **(A)** Phase-locking values across all IN are shown for baseline and three electric field strengths. Individual dots indicate values for each IN in the network. IN that fire at higher intrinsic firing rates (*x* axis) show lower PLV. **(B)** Preferred phases across all IN. At baseline, IN show phase preferences around 0 degree (peak). IN with higher intrinsic firing rates tend to be entrained after the peak. Under tACS, the same trend remained. With increasing electric field strength, IN show larger backward phase shifts.

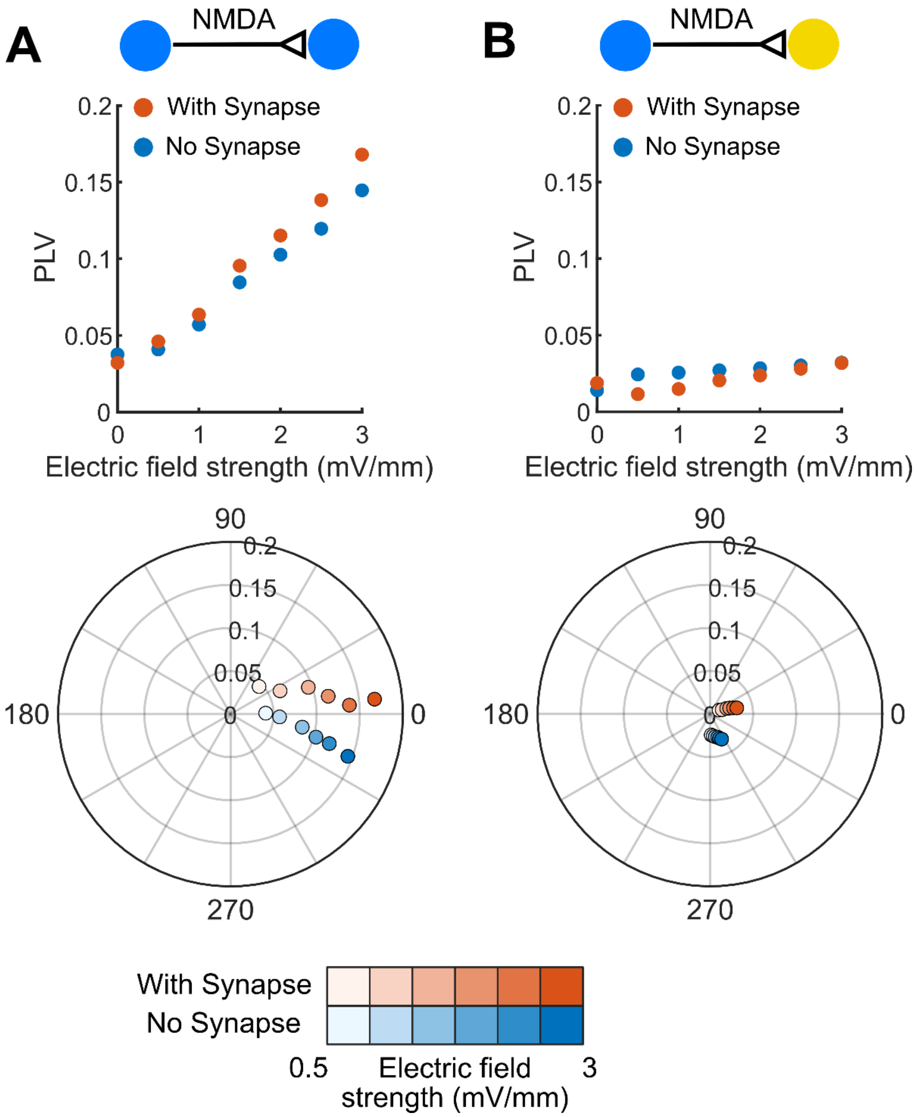


**Supplementary Figure 6. NMDA synapse affects entrainment of the postsynaptic neuron.** Two neurons were connected through a monosynaptic connection. 10 Hz tACS was applied along the somato-dendritic axis of both PY and IN neurons. PLV plots and corresponding polar plots (the radial axis shows PLV) and show the effects of synapse on the entrainment of postsynaptic neuron (with synapse) relative to control (no synapse). **(A, B)** NMDA synaptic connection enhanced entrainment of postsynaptic neurons. The phase preference of the postsynaptic neuron was shifted further backward by NMDA synapses.

**Supplementary Table 1.** Dimensions of neuron compartments (µm)

| | Pyramidal neuron | | Interneuron | |
|---|---|---|---|---|
| | Length | Diameter | Length | Diameter |
| Soma | 30 | 20 | 25 | 25 |
| Dendrite | 2000 | 3.18 | 200 | 3.18 |

**Supplementary Table 2.** Time constants and reversal potentials for each synapse

| Receptor | $\tau_1(ms)$ | $\tau_2$(ms) | $E_{syn}$(mV) |
|---|---|---|---|
| AMPA | 0.2 | 1.7 | 0 |
| NMDA | 2 | 26 | 0 |
| $GABA_A$ | 0.3 | 2.5 | -70 |

**Supplementary Table 3.** Network parameters

| Source | Target | Synapse | Connection Probability | Strength (alpha/low beta) | Delay (ms) |
|---|---|---|---|---|---|
| PY | PY | AMPA | 0.1 | 1.2e-3/1.1e-3 | 1 |
| PY | PY | NMDA | 0.1 | 3e-5/5e-5 | 1 |
| PY | IN | AMPA | 0.1 | 4e-4/4e-4 | 1 |
| PY | IN | NMDA | 0.1 | 3e-5/3e-5 | 1 |
| IN | PY | $GABA_A$ | 0.15 | 1.5e-3/1.5e-3 | 1 |
| IN | IN | $GABA_A$ | 0.1 | 4e-4/4e-4 | 1 |
| Background | PY | Poisson input | 1 | 1.15e-3/1e-3 | 0 |
| Background | IN | Poisson Input | 1 | 6e-5/6e-5 | 0 |
